# BiœmuS: A new tool for neurological disorders studies through real-time emulation and hybridization using biomimetic Spiking Neural Network

**DOI:** 10.1101/2023.09.05.556241

**Authors:** Romain Beaubois, Jérémy Cheslet, Tomoya Duenki, Farad Khoyratee, Pascal Branchereau, Yoshiho Ikeuchi, Timothée Lévi

## Abstract

Characterization and modeling of biological neural networks has emerged as a field driving significant advancements in our understanding of brain function and related pathologies. As of today, pharmacological treatments for neurological disorders remain limited, pushing the exploration of promising alternative approaches such as electroceutics. Recent research in bioelectronics and neuromorphic engineering have led to the design of the new generation of neuroprostheses for brain repair.

However, its complete development requires deeper understanding and expertise in biohybrid interaction. Here, we show a novel real-time, biomimetic, cost-effective and user-friendly neural network for bio-hybrid experiments and real-time emulation. Our system allows investigation and reproduction of biophysically detailed neural network dynamics while promoting cost-efficiency, flexibility and ease of use. We showcase the feasibility of conducting biohybrid experiments using standard biophysical interfaces and various biological cells as well as real-time emulation of complex models. We anticipate our system to be a step towards developing neuromorphic-based neuroprostheses for bioelectrical therapeutics by enabling communication with biological networks on a similar time scale, facilitated by an easy-to-use and accessible embedded real-time system. Our real-time device further enhances its potential for practical applications in biohybrid experiments.

## 1 Introduction

Millions of people worldwide are affected by neurological disorders that strongly impair their cognitive and/or motor functions [1]. An increasing number of technologies and solutions are currently proposed for the treatments of these diseases, whereas being limited to curbing the progress or managing symptoms in most cases [2, 3].

Aside from medical treatment through chemical processes, artificial devices are developed to improve the quality of life of individuals. To bring neuroprosthesis into realization, the behavior of biological neurons as well as its connection and interaction with artificial neural networks must be considered. To this end, investigation of the interaction of neuronal cell assemblies is required to understand and reproduce a specific behavior driven by intrinsic spontaneous activity. Additionally, long-term replacement of damaged brain areas with artificial devices implies understanding of their neurophysiological behaviors.

In this context, new therapeutic approaches and technologies are needed both to promote cell survival and regeneration of local circuits [4] and restore long distance communication between disconnected brain regions and circuits [5]. Thus, characterization and modeling of biological neural networks [6, 7] is crucial to develop new generation of neuroprostheses that mimics biological dynamics and provide adaptive stimulation at biological time scale based on the principle of electroceutics [8, 9].

Thanks to the new neuromorphic platforms, performing bio-hybrid experiments is becoming more and more relevant not only for the development of neuromorphic biomedical devices [8, 9], but also to elucidate the mechanisms of information processing in the nervous system. Recently, major progress has been made in the field of neuroprostheses [6, 7] so as neuromorphic devices are now capable of receiving and processing input while locally or remotely delivering their output either through electrical, chemical or optogenetic stimulation [10].

However, real-time stimulation and processing of biological data using biomimetic Spiking Neural Network (SNN) is still quite rare [11]. Furthermore, to improve temporal accuracy of the stimulation, complex neuron model should be implemented in the SNN [12].

To perform bi-directional bio-hybrid experiments and develop bioelectrical therapeutic solutions for health care like electroceutic [8, 9, 13], real-time biophysics interface and SNN processing are mandatory to ensure interaction at biological time scale [12, 14]. Most of current solutions for biomimetic SNN simulations are software-based such as NEURON [15], NEST [16] or Brian2 [17] tools and show significantly high computation time, especially for complex neuron model with synaptic plasticity. Hence, these latter are not suited for real-time emulation at millisecond time step [18] contrary to hardware-based SNNs. Another benefit of hardware-based SNNs is the ability to perform massive parallel simulations to explore space parameters of neuron models.

In the neuromorphic engineering research, SNNs are designed using two distinct approaches: bioinspired or biomimetic. The former is widely used for applications such as computation and artificial intelligence [19] using accelerated time simulation of simple neuron model. The latter uses complex neuron model operating at biological time scale to simulate neural network dynamics or/and performing bio-hybrid experiments.

Hardware-based SNNs are analog or digital. Analog SNN systems [20] show lower power consumption than digital SNNs [21]. In contrast, digital SNNs are more flexible thus more suited for prototyping while showing overall quicker design time hence constituting the best choice for preliminary experiments and design of new generation of neuroprosthetic. The prominent SNNs hardware platforms are Merolla [22], BrainScaleS-2 [23], SpiNNaker [24] and Loihi [25]. While some of these systems present mobile versions like [26] for BrainScaleS2, they often are not suited for embedded applications. In this manuscript, we present the capabilities of the real-time biomimetic SNN BiœmuS to emulate independent neurons and fully connected networks, showcasing a system integration promoting versatility and ease of use.

## 2 Results

### 2.1 Real-time biomimetic SNN

The low-cost platform targeted is based on a System on Chip (SoC) featuring both Programmable Logic (PL, i.e. FPGA) and processors in a Processing System (PS) part. It is capable of running up to 1,024 neurons fully connected, supporting a total of 2^20^ synapses. It includes on-board monitoring and offers versatile external communication options such as Ethernet, WiFi, expansion PMODs (standard peripheral module interface) and a Raspberry Pi header. The system is used either for real-time emulation as a low-cost computing unit or for biohybrid experiments thanks to its versatility (see Figure 1).

**Fig. 1.**
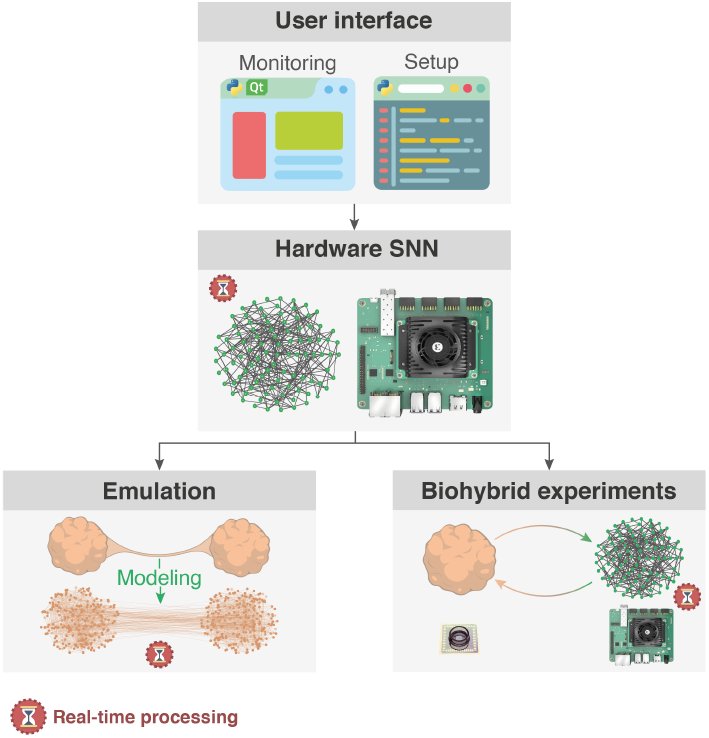
Overview of system applications. The real-time biomimetic SNN implemented in hardware is monitored through a Qt-based GUI and setup by Python scripts ran either on-board or on another computer. The SNN is used either as a real-time emulator for biophysically realistic models or integrated in a biohybrid experiment setup. In a real-time emulation setup, it runs fast simulations of biophysically detailed models suited for large parameters sweeps. Integrated in a biohybrid experimental setup, it acts as a versatile biomimetic artificial neural network easily interfaced with standard biological recording units.

#### 2.1.1 Independents neurons

The neurons composing the SNN are modeled with high biological plausibility using the Hodgkin-Huxley (HH) paradigm [27] in the Pospichil model [28] implementing 6 conductance-based currents. An injected current mimicking synaptic noise following an Ornstein–Uhlenbeck process [29, 30] reproduces spontaneous activity by triggering action potentials on a random basis. All parameters of the HH model as well as the synaptic noise parameters are tuned through the 25 parameters available from the Python scripts (see Figure 2A). The scripts implements 4 preset neuron types including Fast Spiking (FS), Regular Spiking (RS), Intrinsic Burst (IB) and Low Threshold Spiking (LTS) neurons and allow the user to create new presets. The equations of ionic channel states are pre-computed and stored in memory so that they can be easily modified to any channel dynamic without impact on the performances of the system or limitations on mathematical functions used. The computation of ionic currents is performed using 32 bits floating point coding allowing emulation of currents with different dynamics potentially smaller in comparison to other currents like for Ca^2+^-based current in IB or LTS neurons.

**Fig. 2.**
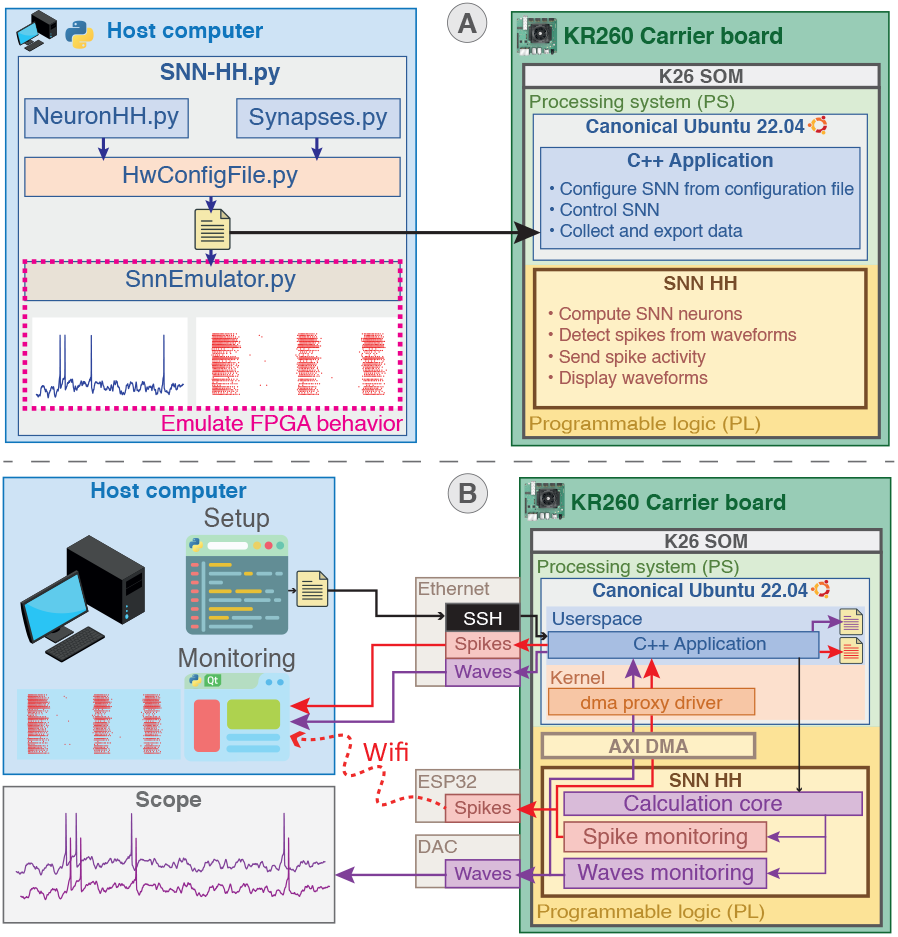
Complete system architecture and integration. **(A)** Overview of system setup from the configuration file generated by Python scripts ran either on-board or on another computer. The configuration file is then read by a C++ application running on Canonical Ubuntu operating system in the Processing System (PS) part to set up the SNN in Programmable Logic (PL) part. Configuration can be emulated beforehand to predict the behavior. **(B)** Schematic of system communication. System control is achieved through the C++ application either remotely *via* SSH or directly on-board from the Ubuntu desktop. Spikes can be monitored concurrently using Ethernet, WIFI and on-board file saving. Waveforms can be monitored concurrently using Ethernet, visualization on scope by probing the Digital-to-Analog Converter (DAC) and on-board file saving.

#### 2.1.2 Connected network

Neurons are connected using biomimetic synapses mimicking AMPA, NMDA, GABA_A_ and GABA_B_ receptors [31] to allow fast and slow synaptic excitation or inhibition, computed using 18 bits fixed point coding. The parameters of the synaptic models can be tuned similarly to the HH parameters through the Python scripts (see Figure 2A). Synaptic connection can be established between all neurons and independently weighted using the Python script allowing the user to create custom functions to setup the connections. The generated configuration file can be emulated using the Python scripts to assess behavior and verify membrane voltage, ionic channel state equations, internal variables and raster plot (see Figure 2A).

#### 2.1.3 Monitoring interface

To maximize compatibility and versatility, a Canonical Ubuntu is running on the processors of the board. Compatibility and versatility are important criteria, knowing that standards for communication protocol interfacing biological recording units vary along with manufacturers (e.g., Serial Peripheral Interface (SPI), Ethernet, USB). In addition, laboratories often have custom setup, designed to reach their specific needs or inherited from prior experimental settings. The selected carrier board features notably multiple USB3.0 and Ethernet ports as well as expansion PMODs (standard peripheral module interface) and Raspberry Pi headers.

The on-board monitoring allows to store all spikes and up to 16 waveforms in a file or/and forward it through ZeroMQ (see Figure 2B). Up to 8 membrane voltage of neurons are selected at a time and output per Digital-to-Analog Converter (DAC) plugged on PMOD connectors. Data is moved from the PL to PS using Direct Memory Access (DMA) interfaced by Advanced eXtensible Interface (AXI) using a driver, thus providing high throughput and good scaling. The interval of collection and forwarding for spikes and waveforms can be set from the application settings.

A wireless setup communication for embedded applications is also provided *via* WiFi using a PMOD ESP32 that plugs on PMOD connectors for spike monitoring. It communicates directly to the PL *via* SPI protocol driven by an ESP32 micro-controller that is able to receive and send data through WiFi network (see Figure 2B). This solution offers a more flexible approach for interconnection of the system that suit well in-vivo applications where cables are a concern, while maintaining a low latency and acceptable throughput. In addition, this constitutes a reusable element to build a reduced and minimal embedded version of the system targeting a smaller programmable logic only target to create an energy-efficient solution for embedded applications.

#### 2.1.4 System control

The SNN is setup from the configuration file generated by Python scripts (see Figure 2A) that is either generated directly on-board using the python installed on the Ubuntu operating system or prior on another computer. The application controlling the system is launched directly using the Ubuntu desktop on the board or remotely over SSH (see Figure 2B). The parameters of the application are generated to JSON format along with the configuration file so as the user may apply changes without code recompilation. The parameters allows to setup the addresses for ZeroMQ forwarding, the local saving or other parameters such as the neurons to monitor. The firmware can be easily updated and loaded by running bash scripts, allowing convenient management of alternative versions developed for a custom dedicated hardware. An external stimulation trigger for each neuron with an independent duration is available *via* ZeroMQ to easily integrate the system in closed-loop setups.

### 2.2 Real-time emulation

This section demonstrates two applications that use BiœmuS as a real-time emulator of biomimetic networks to create a fast emulation setup for large biophysically detailed network.

#### 2.2.1 Interconnected organoids emulation

A more complex network model is emulated representing three-dimensional tissue cultures that are derived from stem cells known as cortical organoids and their interconnections. This model introduces three types of structures promoting different synaptic connections between two organoids as illustrated in Figure 3A.

**Fig. 3.**
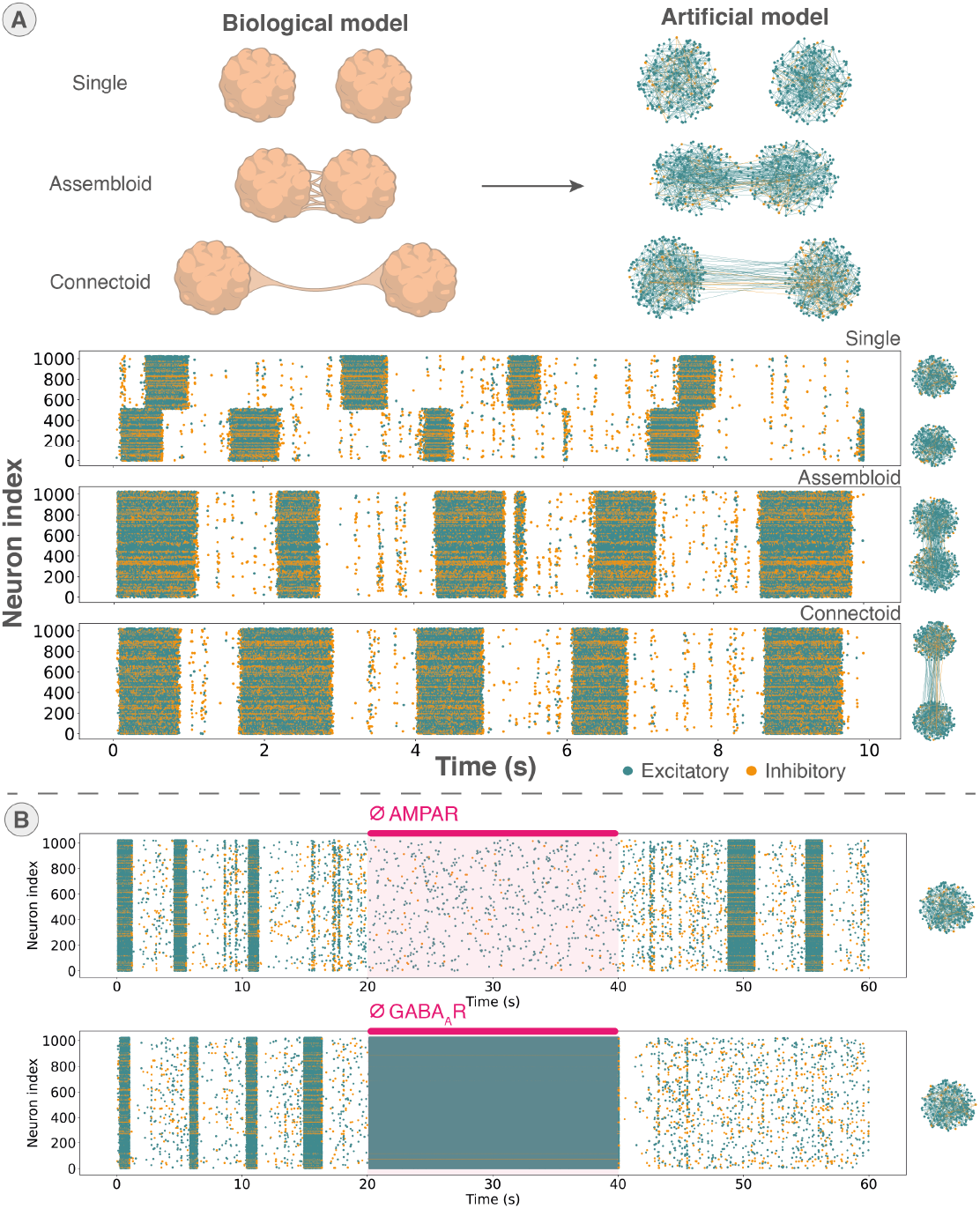
Demonstration applications using BiœmuS. **(A)** Three structures of cortical organoids modeled using FS and RS neurons connected with excitatory and inhibitory synaptic connection (AMPAR and GABA_A_R) based on biological culture observations and their spiking activity. Synaptic connections are promoted according to rules depending on the structure to reproduce, spatial placement of neurons and the ratio of inhibition/excitation connection observed. The spiking activity emulated corresponds to a maximum probability for connection inside and outside the organoids of respectively 10% and 2% with 512 neurons per organoid and a 20% inhibition/excitatory neuron ratio. **(B)** Emulation of drug treatment in a single organoid through AMPAR and GABA_A_R full antagonists from 20 seconds to 40 seconds.

The structure named “single” physically separates the organoids to prevent connection between organoids. It acts as a reference model showing activity of independents organoids. The “assembloid” or fused structure places organoids close to each other thus favouring connection of neurons based on proximity [32]. The “connectoid” structure places organoids centimeters apart while constraining the interconnection to form an axon bundle connecting mostly neurons on the surface of the organoid [33, 34]. The parameters of the SNN were tuned to match the electrical activity in terms of mean firing, synchronicity and burst activity of each structure obtained from MEA recordings.

An additional Python class has been created for that specific model case to assign normally distributed XY coordinates to neurons and generate synaptic connections based on specific rules for each structure. The matrix of connection and list of neurons generated is then simply translated to hardware SNN configuration by the existing software (see Figure 2A), showcasing a case of custom user script to generate the network structure.

The three structures were emulated using 1,024 neurons distributed equally between the two organoids with a similar inhibitory/excitatory ratio to biology. Inhibition is modeled using FS neurons connecting by GABA_A_R and excitation by RS neurons connecting by AMPAR. The emulation is able to reproduce from network bursts to burst synchronization between organoids in the assembloid and connectoid structures as shown in Figure 3A.

#### 2.2.2 Drug treatments emulation

An example of application is the emulation of drug treatments targeting synaptic receptors in an organoid. Two emulations were performed to reproduce a treatment by full antagonist of AMPAR (CNQX) and a treatment by full antagonist to GABA_A_R (Bicuculine). An organoid of similar structure as previously presented is modeled using 1,024 FS and RS neurons connecting with AMPAR and GABA_A_R is emulated on BiœmuS. During emulation, a trigger is sent to BiœmuS to disable a given receptor thus mimicking the drug treatment by full antagonist and a second trigger is sent to reactivate the receptor (see Figure 3B).

We show that the system emulates coherent behavior since the full antagonist to AMPAR prevents bursting and desynchronizes the activity while the full antagonist to GABA_A_R generates continuous spiking activity similar to an epilepsy (see Figure 3B).

### 2.3 Biohybrid experiments

This section presents the biohybrid experiments conducted using the system. It shows how different network implementation from single neuron to larger network can interact with biology through various interfaces.

#### 2.3.1 Open loop biomimetic in-vivo stimulation

A simple case of interaction with the living thanks to the real-time behavior of BiœmuS is to drive open-loop in-vivo stimulation by the SNN [13] as shown in Figure 4A. This open-loop stimulation was applied to rat brains as a neuromorphic-based open-loop set-up for neuroprosthetic applications targeting post-stroke rehabilitation studies [6, 7]. The spikes from neurons emulated by BiœmuS are output as pulses connected to the INTAN RHS recording/stimulation unit to trigger stimulation upon spike reception. The spontaneous activity of the neurons is tuned to obtain slow or fast activities by tuning the parameters of the equation ruling the synaptic noise [13]. In this setup, the latency between spike detection and stimulation is less than a millisecond. This biohybrid experiment promotes the use of BiœmuS as a tool to investigate stroke rehabilitation in an electroceutic approach by providing biomimetic stimulation.

**Fig. 4.**
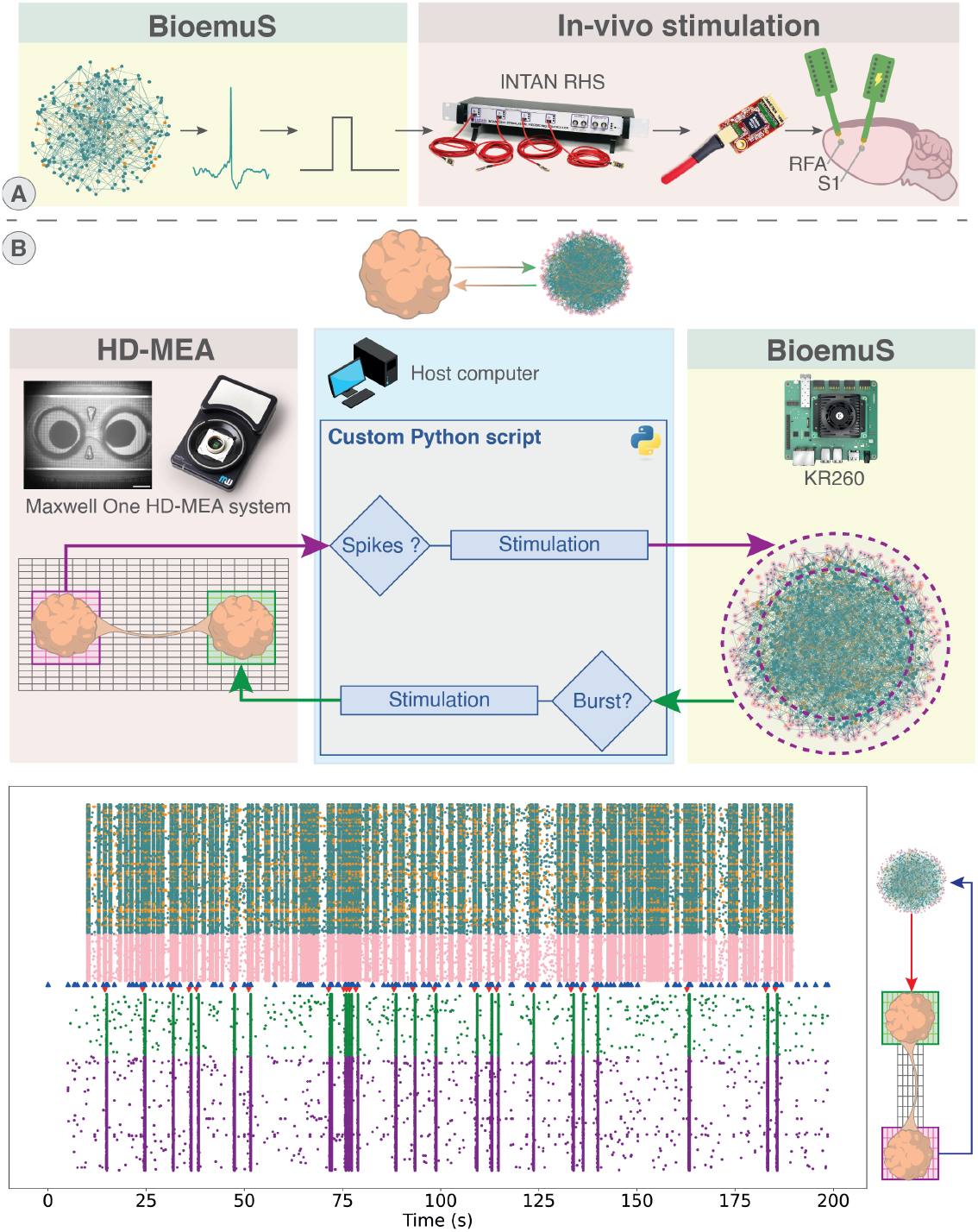
Biohybrid experiments conducted that integrate the system in a biohybrid experimental setup. **(A)** In-vivo stimulation driven by BiœmuS spiking activity as a model of post stroke rehabilitation *via* adaptive stimulation. The spiking activity of the SNN triggers stimulation on an in-vivo culture using the INTAN RHS2116 headstage. Electrode arrays were placed in the rostral forelimb area (RFA) and in the primary somatosensory area (S1) in the brain of adult Long-Evans rats. **(B)** Closed-loop interaction between connected organoids plated on HD-MEA system and single organoid emulated on BiœmuS. The spiking activity detected in the left organoid of the connectoid in the last 100ms triggers stimulation on exterior neurons of the emulated single organoid on BiœmuS. The bursting activity detected on BiœmuS triggers stimulation on the right organoid of the connectoid. Detection and stimulation commands are carried out by Python scripts using. Stimulation on the SNN is performed using the external stimulation slot. BiœmuS stimulation triggers are shown by blue triangle and stimulations to HD-MEA by red triangles. BiœmuS is running for 180 seconds starting from 10 seconds and synchronize manually with HD-MEA activity based on the first stimulation trigger *±* 300 ms.

#### 2.3.2 Closed-loop biomimetic in-vitro stimulation on high resolution MEA

To demonstrate the ease of integration of the system with existing solutions for biological interfacing as well as its versatility, closed-loop stimulation between BiœmuS and the new generation of HD-MEA (High-Density MicroElectrode Array)[35] were performed (see Figure 4B). Connected organoids were plated on HD-MEA. Electrodes were configured to allow activity recording on left and right organoids while allowing stimulation of the right organoid. A single organoid was modeled using BiœmuS on a network of 1,024 neurons and emulating for 180 seconds. Spiking activity of BiœmuS was forwarded to the computer hosting the controlling the HD-MEA system using ZeroMQ over Ethernet and stimulation was sent using ZeroMQ on the external stimulation port of BiœmuS. A Python script executed on that same computer sent stimulation to the HD-MEA upon receipt of a burst from BiœmuS. This experiment showcases the potential of BiœmuS to operate as a tool to study the impact of adaptive stimulation on a culture following the principles of electroceutics while highlighting its ability to adapt to a standard biophysical interface. The benefit of the user-defined model through customizable Python scripts to adapt to a specific application is also showcased here by assigning XY coordinates to neurons to take advantage of the spatial resolution provided by the HD-MEA.

### 2.4 Performances

The low-cost platform targeted is the AMD Xilinx Kria KR260 Robotics Starter Kit carrier board embedding the K26 SOM by AMD Xilinx (Zynq Ultrascale+ MPSoC architecture). This entry level platform is capable of running 1,024 neurons with 6 conductance-based currents for a total of 2^20^ conductance-based synapses running real-time with a time step of 31.25 µs. The system can also run on AMD Xilinx Kria KR260 Vision Starter Kit carrier board with for only restriction the number of PMODs, preventing concurrent from the concurrent use of DAC waveforms and WiFi spike monitoring. While most of the memory available is used, less than 50% of the computing capacity (Logic and Digital Signal Processing slices) of the board is used by the system (see Figure 5). As the design is implemented on an entry level target, the projection of the resources utilization on larger targets suggests the possibility to run several calculation cores in parallel (see Figure 5) as well as allowing faster emulation.

**Fig. 5.**
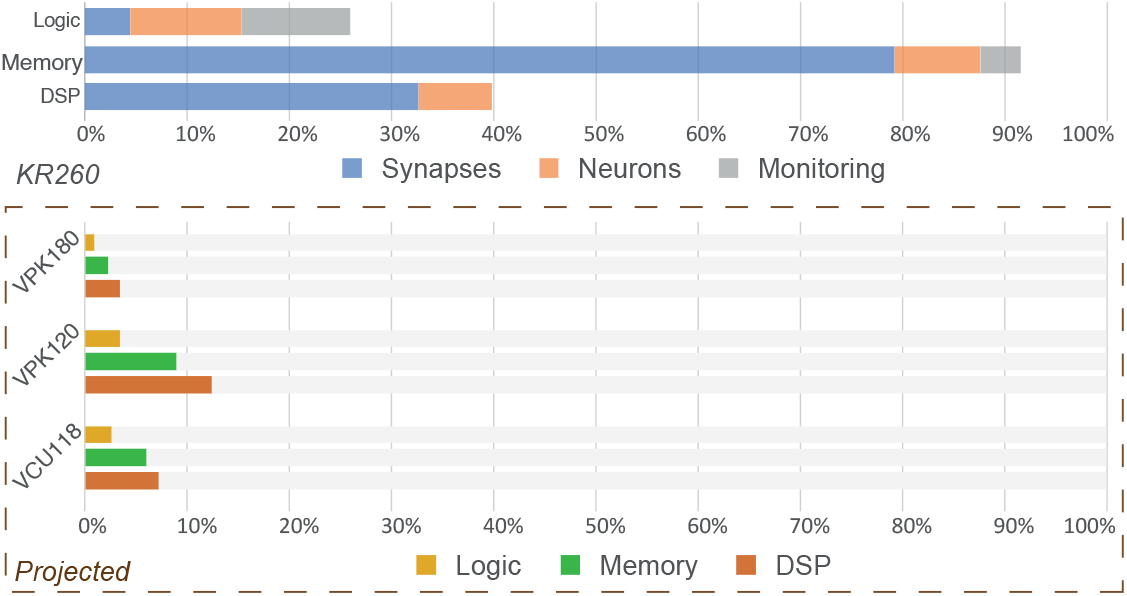
Resources utilization of BiœmuS. Utilization for main modules implemented on AMD Xilinx KR260 Robotic Starter Kit and projected on high end evaluation boards from AMD Xilinx (Versal Premium Series VPK120 and VPK180 Evaluation Kits and Virtex UltraScale+ VCU118 Evaluation Kit). Logic corresponds to LUT and Flip-Flops, memory to the total memory implemented as BRAM and URAM, DSP to the number of Digital Signal Processing (DSP) slices.

The average latency observed to send spikes through Zero MQ (UDP) is 240 µs for 100 ms of spiking activity. The average latency observed for spike monitoring through WiFi (UDP) using ESP32 is between 2.8 ms and 6.2 ms depending on the data collection interval. Overall system power consumption is 6.50W with 3.42W associated with the calculation core. Considering only the calculation core that is running on PL part, BiœmuS consumes 3.42 times more than SpiNNaker [24] or BrainScaleS-2 [23] that run on ASIC.

## 3 Methods

### 3.1 SNN modeling

It uses x ionic channels and mimic better different behavior of a cortical neuron. The synapses model is from [Destexhe et al.] and possesses a biophysical explanation on how synapses work. In addition, a synaptic noise using the ornstein-uhlenbeck process has been used to include spontaneous activities. It has been proven that such models represent the intrinsic noise present in the brain [ref]” something like that.

The neuron model is based on Hodgkin-Huxley [27] in the Pospischil paradigm [28] to guarantee biological meaningfulness while limiting resource consumption and reduce computations. The synapse model used is Destexhe [31] that describes different type of receptors with a conductance-based model that provides biological coherence. Synaptic noise is modeled using Ornstein–Uhlenbeck process that has been proven to represent the intrinsic noise present in the brain [29, 30] that allow the system to create spontaneous activity mimicking biology. The noise seeds are generated by the PS and sent through AXI LITE to the noise generator thus guarantying true random seeds. Equations for ionic channel states are computed from pre-calculated rate stored in memory following the Equation 1 that corresponds to a restated equation of the forward Euler solving.

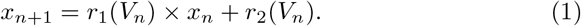

where, *x*_*n*+1_ and *x*_*n*_ are respectively the new and current value of the ionic channel states, *V*_*n*_ is the membrane voltage at previous time step, *r*_1_ and *r*_2_ are the ion rate tables decoded from membrane voltage.

The step and range of the tables are tunable in software but default hardware locks the rate table size to 2048 values (1 BRAM) that provide a good compromise between accuracy and resource usage. The default range is set to −76 mV to 52 mV to provide high accuracy for the preset neurons. Temporal discretization using a small time step compared to the dynamics is chosen to allow explicit numerical solving with forward Euler.

### 3.2 FPGA design

On PL part, the computation core is clocked at 400 MHz, AXI communication to PS at 200 MHz and external components on PMOD connectors such as DAC and ESP32 at 50 MHz. The use of multiple clocks is justified by hardware limitations of components and blocks, multiple clocking allows all parts of the design to work close to their maximum to maximize performances. Crossing clock domain is handled by dual clock BRAM and FIFO for most critical signals, the remaining signals are either handled by double flip-flops or extended. The computation core is fully pipelined.

Computation of ionic channels states and currents are encoded using 32 bits floating point. It grants good stability and accuracy to the computation of ionic channels that are critical parts of the neuron dynamics. Since ionic currents can have different dynamics potentially smaller in comparison to other currents, floating point coding is more suited for most computation and especially for multiplications. Calculation of current sum and forward Euler are encoded using 32 fixed point. Large fixed point coding for sum operations allows to save resources and computation latency compared to floating point, while guarantying consistent accuracy. The synaptic noise, injection current and synapses that have less critical accuracy or perform well with fixed-point coding are computed with 25 and 18 bits fixed point encoding to fit the ranges of DSP slices. Synaptic weight is coded on 14 bits and can be multiplied by a factor specified in software to mimic a larger network behavior.

The numerical solver used is the explicit forward Euler method (Euler–Maruyama) with a small time step compared to the system dynamics to guaranty stability (31.25 µs). To maximize performances and limit resources usage, DSP of the boards were inferred using macros for most operations. The model is validated using Python implementation emulating both rate table based computation and fixed point coding.

### 3.3 System monitoring and control

The PS part is running the Canonical Ubuntu 22.04 for ZynqMP architecture. The main application controlling the SNN is coded and compiled in C++11. Setup from the PS to the PL is implemented by AXI LITE controlled through /dev/mem in the C++ application.

Communication between the PL to PS is implemented using AXI DMA controlled by the the C++ application using the dma proxy driver provided by AMD Xilinx. The application implements a thread for each AXI DMA channel and cyclic buffers for AXI DMA transfers.

The Ethernet communication implements ZeroMQ Push-Pull messaging pattern with a different port for each data (spikes, waveforms, and external stimulation) that can be set from the JSON configuration file.

The interval of data collection can be set from the JSON configuration file from 5ms to 255ms for spike collection *via* DMA, from 3.125 ms to 15 ms for the waveforms collections. The WiFi connection is using UDP protocol and the data collection interval can be set from 2 ms to 20 ms.

The data collection interval for the spikes and waveforms through the DMA directly impacts the load of the application. A small interval will generate more frequent write in file or frame sending thus loading the CPU. The limit corresponds to a data collection interval smaller than the writing or sending time of the frame therefore blocking the software in a thread.

The data collection interval for WiFi forwarding is limited by the hardware and latency of the WiFi protocol so as high interval generates too large buffer and too small interval may generate packet loss.

DMA based monitoring can run local saving and Ethernet forwarding concurrently in most cases with large data collections interval but may dysfunction on small interval due to processor performances. Spikes and waveforms monitoring through DMA can run concurrently in separate threads but may also dysfunction on small data collection intervals due to processor performances. WiFi, DAC and DMA based monitoring can run concurrently without impact on performances. Bash scripts are used to compile the software, update the firmware and launch the application.

An external stimulation controlled *via* Ethernet over ZeroMQ allows to send a stimulation of a given time to a given neuron by passing the stimulation duration and neuron index to the PL using the AXI DMA.

### 3.4 Real-time emulation

#### Interconnected organoids emulation

The “single” physically physically separates the organoids to prevent connection. The “assembloid” or fused places organoids tens of micrometers apart [32]. The “connectoid” places organoids centimeters apart while constraining the interconnection to a channel of 150 µm width [33, 34]. The emulation model implements cortical neurons using FS and RS types connected by AMPAR and GABA_A_R.

The synaptic connection rules for the synaptic connections inside organoids are ruled by Equation 2 that favors connection to neurons close to each other normalised by the diameter of organoid. The connections between organoids are ruled by Equation 3 for assembloid and by Equation 4 for connectoid. The former favors connection to neurons close to each other normalised by the maximum distance possible between neurons, while the connectoid rule is promoting connection based on the location of neuron in the organoid that promots connection on the exterior ring.

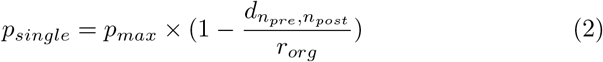

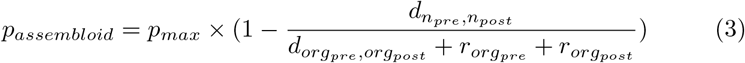

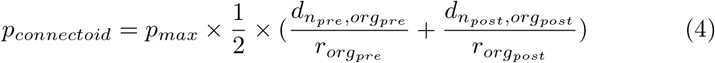

where *p*_*max*_ is the maximum probability of connection, *d* is the distance, *diam*_*org*_ the diameter of the organoid, *r* the radius, *n*_*pre*_ and *n*_*post*_ the presynaptic and post-synaptic neurons, *org*_*pre*_ and *org*_*post*_ the pre-synaptic and post-synaptic organoids and the distance calculated from the center of the organoids.

#### Drug treatment emulation

The organoid emulated corresponds to 1,024 neurons distributed in 10 % of FS neurons and 90 % of RS neurons. FS neurons connect with GABA_A_R while RS neurons connect with AMPAR. The synaptic connections inside the organoids were generated using the same algorithm as for the single structure (Equation 2). The control of the activation and inactivation of the synapses is handled by an AXI LITE register that was set from an external computer using the same port as external stimulation trigger (Ethernet over ZeroMQ). The python script sending the trigger from the external computer was designed to disable synaptic connections of BiœS after 20 seconds of emulation and reactivate after 20 seconds. The python was synchronized by using a blocking call on the availability of BiœmuS to receive frames as it becomes available only after the emulation started. For the full antagonist AMPAR, the AMPA calculation block was disabled and the GABA_A_R in the case of the full antagonist GABA_A_R. The activation and inactivation of the synapses is done by conditional consideration of the synaptic current in the sum. The spiking activity was recorded using the on-board saving of spikes with a data collection interval of 100 ms.

### 3.5 Biohybrid experiments

#### Open-loop biomimetic in-vivo stimulation

The experiment shown in Figure 4A corresponds to a former version of BiœmuS implementing only independent neurons using exclusively fixed point coding and fitted equations for ionic channel states based on [36]. The platform was the ZyboZ7-20 running the C++ application in standalone mode with spike monitoring polled using AXI LITE and forwarded to the host computer through USB 2.0 CDC. The parameters of the FS and RS neurons used are the same as in [36]. A spike was considered in hardware when the membrane potential of a neuron crossed −10 mV and generated a pulse on a 3.3V digital output. The experiment conducted corresponds to the work [13] that provides further details on the experimental setup and protocol.

Healthy adult Long-Evans rats (5 male, weight: 300-400g, age: 4-5 months; Charles River Laboratories, Calco, LC, Italy) were employed for this work. All the rats were treated with the SNN-based stimulation while they were deeply anesthetized. The experimental procedures were performed in the Animal Facility of the Italian Institute of Technology (IIT), Genoa, Italy and were previously approved by the Italian Ministry of Health and Animal Care (Italy: authorization n. 509/2020-PR).

Anesthesia was induced by placing the rat inside a vaporizing chamber and injecting gaseous isoflurane (5% @ 1 lpm). The surgical level of anesthesia was induced by the administration of ketamine (80-100 mg/kg IP) and xylazine (5-10 mg/kg). The rat was then secured in a stereotaxic frame and all vital parameters were monitored until the end of the procedure. The surgery began by applying lidocaine cream (a topical analgesic) before performing a midline skin incision to expose the skull. Successfully, a laminectomy was performed at the level of the Cisterna Magna to allow the draining of cerebrospinal fluid (CSF). Then, based on stereotaxic measurements [9] +3.5, +2.5 and –1.25, +4.25 AP, ML, burr holes (3 mm diameter) were performed over the primary somatosensory area (S1) and rostral forelimb area (RFA). Lastly, the dura mater was removed from the burr holes (RFA and S1) to allow insertion of MEAs (MEAs; A4×4-5 mm-100-125-703-A16, NeuroNexus)

#### Closed-loop biomimetically driven stimulation on HD-MEA

The bidirectional communication between BiœmuS and the HD-MEA system is ensured by Python scripts running on a gateway computer. The HD-MEA was configured to record from channels both from left and right organoid based on an activity scan and to select random stimulation electrodes on the right organoids. The HD-MEA is the MaxOne chip of MaxWell Biosystems AG. The spikes received from BiœmuS on the host computer are analyzed to detect the presence of a burst in the 100 ms of activity sent. A burst is defined as more than 64 neurons spiking at least 15 times in the last 100ms. Upon burst detection, a stimulation of one period of a 100Hz sinus wave with an amplitude of 40 mV is sent to the HD-MEA using custom Python script based on manufacturer templates. Stimulation was chosen of amplitude high enough to allow visualization of the stimulation on the MaxLab Live Software.

The spikes received from the HD-MEA triggered stimulation on BiœmuS if at least 1 spike was detected on at least 2 channels in the last 100ms of activity collected. The stimulation was sent through Ethernet over ZeroMQ to the external stimulation port of BiœmuS to trigger a stimulation of 6.250ms of 0.03 *mA/cm*^2^ on the neurons on the exterior rings of the organoid.

The Python script implemented executed a thread for each task of receiving spikes from HD-MEA, receiving spikes from BiœmuS, sending stimulation to Maxwell and sending stimulation to BiœmuS.

The activity of the HD-MEA was recording using the MaxLab Live Software started manually before starting BiœmuS. The activity was analysed using the script provided by the manufacturer. The spiking of activity of BiœmuS was recorded on-board.

The configuration of electrodes of the HD-MEA was exported from the software. The XY configuration of neurons, network configuration and stimulated neurons of BiœmuS were exported from the Python scripts. Detection of burst and spikes triggering stimulation for both HD-MEA and BiœmuS were reconstructed from the recorded data. The synchronization of both activities was done manually based on the trigger of the first stimulation considering an approximation of 100 to 300 ms based on the latency of the HD-MEA communication and the fluctuating latency induced by the Ubuntu operating system.

#### Organoid cultures

Cortical connectoids were generated using previously reported protocol [37]. Briefly, hiPSCs were dissociated using TrypLE Express and 10,000 cells per well were seeded into U-bottom ultra-low attachment 96 well plate (Prime surface, Sumitomo bakelite) in mTeSR plus supplemented with 10µM of Y-23632. 24h later, media was replaced with neural induction media (NIM), consisting of DMEM-F12 with HEPES, 15% (v/v) knockout serum replacement, 1% (v/v) minimal essential media non-essential amino acids (MEM-NEAA), and 1% (v/v) Glutamax, supplemented with 100 nM LDN-193189, 10 µM SB431542, and 5% (v/v) heat-inactivated FBS. On day 2, NIM was replaced without the supplement of FBS and changed every other day until day 10.

From day 10 to 18, culture medium was replaced and changed every other day with neural differentiation media 1 (NDM1), consisting of 1:1 mixture of DMEM/F12 with HEPES and Neurobasal medium, 0.5% (v/v) N2 supplement, 1% (v/v) B27 supplement without vitamin A, 1% (v/v) Glutamax, 0.5% (v/v) MEM-NEAA, 0.25 mg/ml human insulin solution, and 1% (v/v) Penicillin/Streptomycin/Amphotericin (PSA) (Sigma, A5955). On day 18, culture medium was replaced with neural differentiation media 2(NDM2), consisting of Neurobasal medium, 0.5% (v/v) N2 supplement, 1% (v/v) B27 supplement with vitamin A, 1% (v/v) Glutamax, 0.5% (v/v) MEM-NEAA, 0.25 mg/ml human insulin solution, 200 mM ascorbic acid, and 1% (v/v) PSA, supplemented with 20 ng/ml brain derived neurotrophic factor (BDNF). On day 28, culture media was replaced with Neural Maintenance Media (NMM) consisting of Neurobasal Medium, supplemented with 2% (v/v) B27 supplement with vitamin A, 1% (v/v) Glutamax, 1% (v/v) PSA and 20 ng/ml BDNF.

Cerebral organoids were subjected to connectoid formation after 60 days in culture. Here, a costume made microfluidic device containing two holes which are connected through a narrow channel were bonded on a CMOS-based HDMEA (MaxOne, Maxwell Biosystems). Microchannel of the microfluidic device was coated with 2% Matrigel (Corning) in DMEM/F12 for 1h at room temperature (RT). Next, coating solution is replaced with NMM and an organoid is placed into each of the holes. Cells were kept at 37°C and 5% CO2 and half media change was performed every 3-4 days for the duration of cell culture.

## 4 Discussion

Not applicable.

## 5 Conclusion

Running a generic operating system on the PS to handle communication offers versatility and ease integration with existing experimental setups, while reducing development time where the low-level FPGA development is technical and time consuming. Another benefit is the ease of use for biologists thanks to the graphic interface and user-friendly approach offered by an Ubuntu operating system. While non real-time operating system as Ubuntu induces a discernible and fluctuating latency, using PL driven interrupt and AXI DMA allows to obtain relatively low latency about the tens of microseconds. A trade-off between latency and compatibility/versatility can be found by using solutions such as data sent directly by PL trough expansion PMODs or ESP32, realtime operating system or running the application the real-time cores of the chip. Nonetheless, direct monitoring on the PL that drastically reduces the latency remains possible using the various connectors of the board but at the cost of longer and more complex development.

On the current target, the main bottleneck lies in the memory usage essentially allocated for synapses weights and pre-calculated ionic channel states. Since the current target is using a preceding architecture, more efficient architectures of memory can be found in recent larger targets such as High Bandwidth Memory (HBM) that integrates DRAM directly into the FPGA package, thus providing drastically higher depth and bandwidth. Latest AMD Xilinx chips also incorporate adaptive SoCs that provide significantly higher computation power notably with native floating point DSP and AI engine while still embedding a Zynq for setup and control Figure 5. Hence porting a similar architecture of SNN on these targets would significantly increase performances and create a *via*ble alternative to standard GPU. An alternative would be to reduce the number of synapses as fully connected network is not always necessary, thus allowing the implementation of more neurons.

The system has proven its ease of integration demonstrated by the biohybrid experiments conducted on most widespread biophysical interface where low-level communication protocol (pulse on digital output) as well as complex communication protocols (WiFi and Ethernet) were implemented. The ease of use also has been particularly promoted by the application Figure 3A showing an example of complex network could be created simply from a customizable Python script. The experiment in Figure 4B also highlighted this feature by interfacing the BiœmuS to a biophysical interface using only Python scripts.

The presented applications demonstrate the flexibility of BiœmuS in adapting to the study of various biological processes, including stroke trough in-vivo stimulation (see Figure 4A) and the potential for neuroprostheses replacement through closed-loop in-vitro stimulation driven by BiœmuS (see Figure 4B).

We are proposing a low-cost, flexible and real-time biomimetic tool that could allow wider exploration of the mechanism of the living thanks to realtime emulation and hybridization.

## Supplementary information

Not applicable.

## Acknowledgments

We acknowledge Landry Bailly for the synaptic connection heatmap of the organoid emulation, Andréa Combette for the analysis of the spiking activity of the organoid emulation, Mattia Di Florio and Marta Carè for the preparation and execution of the experiment on open-loop in-vivo stimulation. We also acknowledge Ryota Murai for his advice on the RedPitaya system, Hu Huaruo and Atsuhiro Nabeta for their advice and assistance on the data acquisition from Maxwell system.

## Declarations

### Ethics approval

Not applicable.

### Consent to participate

Not applicable.

### Consent for publication

Not applicable.

### Availability of data and materials

The data analysed in this study are available from the corresponding authors upon reasonable request.

### Code availability

The code related to the experiments and tests is available from the corresponding authors upon reasonable request.

### Author contributions

R.B. designed both the software and hardware part of the system, developed the Python scripts, performed the biohybrid experiments, analysed the results and wrote the manuscript. J.C. participated in the development of the hardware design of the synapses, WiFi communication on ESP32 and designed the reduced version controlling snake robot. T.D. cultivated the organoids, performed the analysis of the data from the Maxwell system and captured the images of the cultures. F.K. participated in the design of the reduced version working controlling the snake robot. T.L. supervised and participated in the design of the applications and biohybrid experiments. Y.I. and P.B. supervised and advised on the biohybrid experiments and biological modeling. T.L., Y.I. and P.B joined the discussion and corrected the draft manuscript. All authors discussed and revised the final manuscript.

## Appendix A Section title of first appendix

## References

[1] Organization, W.H., et al.: The top 10 causes of death; 24 May 2018 (2020)

[2] Chin, J.H., Vora, N.: The global burden of neurologic diseases. Neurology 83(4), 349–351 (2014)

[3] French, B., Thomas, L.H., Coupe, J., McMahon, N.E., Connell, L., Harrison, J., Sutton, C.J., Tishkovskaya, S., Watkins, C.L.: Repetitive task training for improving functional ability after stroke. Cochrane database of systematic reviews (11) (2016)

[4] Farina, D., Vujaklija, I., Brånemark, R., Bull, A.M., Dietl, H., Graimann, B., Hargrove, L.J., Hoffmann, K.-P., Huang, H., Ingvarsson, T., et al.: Toward higher-performance bionic limbs for wider clinical use. Nature biomedical engineering, 1–13 (2021)

[5] Bouton, C.E., Shaikhouni, A., Annetta, N.V., Bockbrader, M.A., Friedenberg, D.A., Nielson, D.M., Sharma, G., Sederberg, P.B., Glenn, B.C., Mysiw, W.J., et al.: Restoring cortical control of functional movement in a human with quadriplegia. Nature 533(7602), 247–250 (2016)

[6] Panuccio, G., Semprini, M., Natale, L., Buccelli, S., Colombi, I., Chiappalone, M.: Progress in neuroengineering for brain repair: New challenges and open issues. Brain and neuroscience advances 2, 2398212818776475 (2018)

[7] Semprini, M., Laffranchi, M., Sanguineti, V., Avanzino, L., De Icco, R., De Michieli, L., Chiappalone, M.: Technological approaches for neurorehabilitation: from robotic devices to brain stimulation and beyond. Frontiers in neurology 9, 212 (2018)

[8] Famm, K., Litt, B., Tracey, K.J., Boyden, E.S., Slaoui, M.: A jump-start for electroceuticals. Nature 496(7444), 159–161 (2013)

[9] Reardon, S., et al.: Electroceuticals spark interest. Nature 511(7507), 18 (2014)

[10] Christensen, D.V., Dittmann, R., Linares-Barranco, B., Sebastian, A., Le Gallo, M., Redaelli, A., Slesazeck, S., Mikolajick, T., Spiga, S., Menzel, S., et al.: 2022 roadmap on neuromorphic computing and engineering. Neuromorphic Computing and Engineering 2(2), 022501 (2022)

[11] Xu, T., Xiao, N., Zhai, X., Chan, P.K., Tin, C.: Real-time cerebellar neuroprosthetic system based on a spiking neural network model of motor learning. Journal of Neural Engineering 15(1), 016021 (2018)

[12] Sharifshazileh, M., Burelo, K., Sarnthein, J., Indiveri, G.: An electronic neuromorphic system for real-time detection of high frequency oscillations (hfo) in intracranial eeg. Nature communications 12(1), 3095 (2021)

[13] Di Florio, M., Caré, M., Beaubois, R., Barban, F., Levi, T., Chiappalone, M.: Design of an experimental setup for delivering intracortical microstimulation in vivo via spiking neural network. In: 2023 45th Annual International Conference of the IEEE Engineering in Medicine & Biology Society (EMBC) (2023). IEEE

[14] Corradi, F., Indiveri, G.: A neuromorphic event-based neural recording system for smart brain-machine-interfaces. IEEE transactions on biomedical circuits and systems 9(5), 699–709 (2015)

[15] Hines, M.L., Carnevale, N.T.: Neuron: a tool for neuroscientists. The neuroscientist 7(2), 123–135 (2001)

[16] Gewaltig, M.-O., Diesmann, M.: Nest (neural simulation tool). Scholarpedia 2(4), 1430 (2007)

[17] Stimberg, M., Brette, R., Goodman, D.F.: Brian 2, an intuitive and efficient neural simulator. Elife 8, 47314 (2019)

[18] Van Albada, S.J., Rowley, A.G., Senk, J., Hopkins, M., Schmidt, M., Stokes, A.B., Lester, D.R., Diesmann, M., Furber, S.B.: Performance comparison of the digital neuromorphic hardware spinnaker and the neural network simulation software nest for a full-scale cortical microcircuit model. Frontiers in neuroscience 12, 291 (2018)

[19] Tavanaei, A., Ghodrati, M., Kheradpisheh, S.R., Masquelier, T., Maida, A.: Deep learning in spiking neural networks. Neural networks 111, 47–63 (2019)

[20] Donati, E., Payvand, M., Risi, N., Krause, R., Indiveri, G.: Discrimination of emg signals using a neuromorphic implementation of a spiking neural network. IEEE transactions on biomedical circuits and systems 13(5), 795–803 (2019)

[21] Davidson, S., Furber, S.B.: Comparison of artificial and spiking neural networks on digital hardware. Frontiers in Neuroscience 15, 651141 (2021)

[22] Merolla, P., Arthur, J.V., Alvarez-Icaza, R., Cassidy, A.S., Sawada, J., Akopyan, F., Jackson, B.L., Esser, S.K., Appuswamy, R., Taba, B., Amir, A., Flickner, M.: Merolla communication network and interface a million spiking-neuron integrated circuit with a scalable. (2014)

[23] Pehle, C., Billaudelle, S., Cramer, B., Kaiser, J., Schreiber, K., Stradmann, Y., Weis, J., Leibfried, A., Müller, E., Schemmel, J.: The brainscales-2 accelerated neuromorphic system with hybrid plasticity. Frontiers in Neuroscience 16 (2022)

[24] Painkras, E., Plana, L.A., Garside, J., Temple, S., Galluppi, F., Patterson, C., Lester, D.R., Brown, A.D., Furber, S.B.: Spinnaker: A 1-w 18-core system-on-chip for massively-parallel neural network simulation. IEEE Journal of Solid-State Circuits 48(8), 1943–1953 (2013)

[25] Davies, M., Srinivasa, N., Lin, T.-H., Chinya, G., Cao, Y., Choday, S.H., Dimou, G., Joshi, P., Imam, N., Jain, S., et al.: Loihi: A neuromorphic manycore processor with on-chip learning. Ieee Micro 38(1), 82–99 (2018)

[26] Stradmann, Y., Billaudelle, S., Breitwieser, O., Ebert, F.L., Emmel, A., Husmann, D., Ilmberger, J., Müller, E., Spilger, P., Weis, J., et al.: Demonstrating analog inference on the brainscales-2 mobile system. IEEE Open Journal of Circuits and Systems 3, 252–262 (2022)

[27] Hodgkin, A., Huxley, A.: A quantitative description of membrane current and its application to conduction and excitation in nerve. Bulletin of Mathematical Biology 52(1-2), 25–71 (1990). 10.1016/s0092-8240(05)80004-7

[28] Pospischil, M., Toledo-Rodriguez, M., Monier, C., Piwkowska, Z., Bal, T., Frégnac, Y., Markram, H., Destexhe, A.: Minimal Hodgkin-Huxley type models for different classes of cortical and thalamic neurons. Biological Cybernetics 99(4-5), 427–441 (2008). 10.1007/s00422-008-0263-8

[29] Destexhe, A., Rudolph, M., Fellous, J.M., Sejnowski, T.J.: Fluctuating synaptic conductances recreate in vivo-like activity in neocortical neurons. Neuroscience 107(1), 13–24 (2001). 10.1016/S0306-4522(01)00344-X

[30] Grassia, F., Kohno, T., Levi, T.: Digital hardware implementation of a stochastic two-dimensional neuron model. Journal of Physiology Paris 110(4), 409–416 (2016). 10.1016/j.jphysparis.2017.02.002

[31] Destexhe, A., Mainen, Z.F., Sejnowski, T.J.: Kinetic models of synaptic transmission: From Ions to Networks. Methods in Neural Modeling: from Ions to Networks, 1–25 (1998)

[32] Pąsca, S.P.: Assembling human brain organoids. Science 363(6423), 126–127 (2019)

[33] Kirihara, T., Luo, Z., Chow, S.Y.A., Misawa, R., Kawada, J., Shibata, S., Khoyratee, F., Vollette, C.A., Volz, V., Levi, T., et al.: A human induced pluripotent stem cell-derived tissue model of a cerebral tract connecting two cortical regions. Iscience 14, 301–311 (2019)

[34] Kawada, J., Kaneda, S., Kirihara, T., Maroof, A., Levi, T., Eggan, K., Fujii, T., Ikeuchi, Y.: Generation of a motor nerve organoid with human stem cell-derived neurons. Stem cell reports 9(5), 1441–1449 (2017)

[35] Ballini, M., Müller, J., Livi, P., Chen, Y., Frey, U., Stettler, A., Shadmani, A., Viswam, V., Jones, I.L., Jäckel, D., et al.: A 1024-channel cmos microelectrode array with 26,400 electrodes for recording and stimulation of electrogenic cells in vitro. IEEE journal of solid-state circuits 49(11), 2705–2719 (2014)

[36] Khoyratee, F., Grassia, F., Säighi, S., Levi, T.: Optimized real-time biomimetic neural network on fpga for bio-hybridization. Frontiers in neuroscience 13, 377 (2019)

[37] Osaki, T., Ikeuchi, Y.: Advanced complexity and plasticity of neural activity in reciprocally connected human cerebral organoids. BioRxiv, 2021–02 (2021)

